# Dissecting the landscape of activated CMV-stimulated CD4+ T cells in human by linking single-cell RNA-seq with T-cell receptor sequencing

**DOI:** 10.1101/2020.08.30.268433

**Authors:** Menghua Lyu, Shiyu Wang, Kai Gao, Longlong Wang, Bin Li, Lei Tian

**Affiliations:** BGI Education Center, University of Chinese Academy of Sciences, Shenzhen 518083, China; BGI-Shenzhen, Shenzhen, China 518083; Shanghai Institute of Immunology, Shanghai JiaoTong University School of Medicine, Shanghai 200025, China

**Author notes:** These authors contributed equally to this work (Menghua Lyu, Shiyu Wang).

**Keywords:** CMV pp65, Single-cell mRNA-seq, paired TCR-seq, CD4+ T cells

## Abstract

CD4 T cell is crucial in CMV infection, but its role is still unclear during this process. Here, we present a single-cell RNA-seq together with T cell receptor (TCR) sequencing to screen the heterogenicity and potential function of CMV pp65 reactivated CD4+ T cell subsets from human peripheral blood, and unveil their potential interactions. Notably, Treg composed the major part of these reactivated cells. Treg gene expression data revealed multiple transcripts of both inflammatory and inhibitory functions. Additionally, we describe the detailed phenotypes of CMV-reactivated effector-memory (Tem), cytotoxic T (CTL), and naïve T cells at the single-cell resolution, and implied the direct derivation of CTL from naïve CD4+ T cells. By analyzing the TCR repertoire, we identified a clonality in stimulated Tem and CTLs, and a tight relationship of Tem and CTL showing a large share in TCR. This study provides clues for understanding the function of CD4+ T cells subsets and unveils their interaction in CMV infection, and may promote the development of CMV immunotherapy.

## Introduction

Cytomegaloviruses (CMV)/human herpesvirus 5 (HHV-5) infection is endemic in humans. Most immunocompetent hosts show little to no clinical symptoms of primary infection and during persistent infection. Although its infection is regarded as asymptomatic, CMV hijacks the resources of the immune system throughout life by remaining latent and occasionally reactivate, eventually compromising on average approximately 10% of the entire T cell repertoire^1^ and had a deleterious effect on immune senescence and health outcomes in the elderly^2^. In addition to this impact in healthy people, CMV infection can cause devastating consequences on immuno-compromised populations, such as the fetus and patients undergoing transplantation.

For these immunocompromised patients, reconstruction of CMV-specific T cells has emerged as an effective method to reduce CMV infection and reactivation. Data from patients who have received hematopoietic stem cell transplantations (HSCT) shows that the recovery from CMV disease correlates with the reconstitution of CD4+ and CD8+ T cell pools ^3–5^, and the recovery of CD4+ T cells is suggested as a prerequisite. The underlying mechanism may be that CMV-specific CD4+ T cells affiliate the expansion of CMV-specific CD8+ T cells, and lead a more effective clearance of serum virus compared to treatment with CD8+ T cells only^6^. Furthermore, infusion experiments with CD4+ T cells alone in immunocompromised mice is able to effectively repress CMV reactivation. These evidences suggest the key effector of CD4+ T cells in anti-CMV immunity, but CD4+ T is a heterogeneous group, and it is yet unclear of the function of these CD4+ T subsets in CMV infection and their interaction which hesitate the clinical application of adoptive immune therapy in CMV.

Therefore, studies on CD4+ T cell subsets have been performed respectively in past decades and reveal that CD4+ cytolytic cell (CD4-CTL), Treg and CD4+ memory T involve in the immune response to CMV infection in humans, nonhuman primates, and rodents. CD4-CTLs are firstly confirmed as a natural identity in chronic infections, such as LCMV, HBV, and CMV, and shows a strong antiviral effect in anti-CMV immunity by their helper functions and cytotoxicity. In terms of its helper function, CD4-CTL expressed cytokines, such as IFNγ and TNF^7^, to promote the activation of CD8+ T cells, recruit innate immune cells including natural killer and monocytes to inflammatory sites, and directly inhibit virus replication^8^. As for their cytotoxicity, Fas/FasL pathway is utilized to mediate the death of infected B cells presenting viral epitopes with MHC-II^9,10^. Another cytotoxic mechanism used by CD4-CTL is via the perforin–granzyme pathway^11^. This pathway is based on the recognition of CTL to target cells in an MHC-II dependent manner^12^, where the MHC-II is upregulated in epithelial cells upon CMV infection. Although there have been advances in understanding CD4-CTL function in CMV infection, their derivation is still unclear. Common views developed from other infection diseases indicate the origination of CD4-CTLs from effector cells. Recently, evidences^13,14,15^ from studies on transcriptome factors suggested these cells can also differentiate from naïve cells directly. This indicates that it may be possible to prepare CD4-CTL for immune therapy from naïve cells.

Although some studies tried to unveil Treg function in CMV infection, it is still controversial to conclude. In humans, Treg cells from CMV-seropositive individuals attenuated the proliferation of autologous CD8+ T cells and, to a lesser extent, CD4+ T cells in response to CMV virus ex vivo stimulation using PD-1 pathway^16^. However, in CMV reactivating patients received HSCT, CMV reactivation does not correlate with the numerical reconstitution of CD4+CD25highCD127-Tregs, and conventional T cells in these patients expressed high level of Ki67 indicating that their activation function is unimpaired^17^. Selectively removing Treg in animal models is a classical method to verify Treg function at infectious situations^18^, and have been used to identify the negative function of Treg in some anti-viral immunities. However, these experiments failed to conclude Treg function in CMV infection. In murine, eight months after infection, Treg cells deletion decreased MCMV reaction in the spleen but enhance the reactivation in the salivary gland ^19^. By analyzing the cell composition and cytokine secretion, the Treg deletion was found to expanded CD4+ T cells in these two tissues, excepting of an increased production of IL-10 in the salivary gland which is inferred to disturb the conventional T cells function in the salivary gland and be in charge of the enhancing viral replication.

T cell receptor (TCR)-specific signaling pathway is essential to generate an effective antiviral immunity. To identify T cells bearing these antigen-specific TCRs, labeling them with antibodies targeting IFN-γ and other markers, such as CD69, as well as Fluorescence-activated cell sorting (FACS) are used in previous studies. In the past decade, CD154 is found to be a TCR-singling specific marker, and labeling it only is able to identify cells which should be labeling with multiple markers^20–22^. Therefore, methods based on CD154 have been developed and employed by studies focusing on the antigen specificity of TCRs^23,24^

To further unveil the potential function of CD4+ T cell subsets in CMV and understand their interactions, we performed a single-cell RNA and paired TCR sequencing on CMV pp65-specific CD4+ T cells from three healthy donors with a latent CMV infection. With a global view on CMV-specific CD4+ cells, we identified: 1) CMV-reactivated Treg cells accounted of a large proportion, and obtained a Th1 phenotype, enhanced migration ability and multiple inhibitory functions; 2) CD4-CTLs have a polyfunctional phenotypes; 3) CD4-CTL and effector memory T (Tem) experienced clonality and have a lager convergence in TCR repertoire; 4) a group of naïve cells expressed cytolytic factor. These findings exhibited the heterogeneity of CMV-reactivated CD4+ T cells, highlighted the balance between CMV-specific Treg and effector T cells, suggest that the composition of CD4+ T cells may be critical for adoptive T cells therapy for CMV. In summary, this study provides clues for understanding anti-CMV immunity and developing clinical therapy on CMV.

## Results

### CMV pp65-specific CD4+ T cells are characterized by typical antiviral profiles

Circulating antigen-specific T cells are rare in peripheral blood during the latent stage of CMV infection, comprising on average 10% of both the CD4 and CD8 memory compartments in blood^1^. To isolate CMV-specific CD4T cells, we cultured PBMCs with and without CMV-pp65 peptides for 24 hours and sorted CD3+CD154+ cells by flow cytometry^24–27^. We sorted CMV-specific CD4T cells from three donors and then pooled them together for single-cell mRNA-seq and paired VDJ-seq using the 10 × Chromium platform (Figure 1A). Control CD4T cells were acquired by lymphocyte sorting with FSC/SSC. PBMCs from the three donors were subject to bulk RNA-seq for subsequent SNP (single-nucleotide polymorphism) calling, and the sample identity of each cell was deconvoluted with these natural genetic variations^28^.

**Figure 1:**
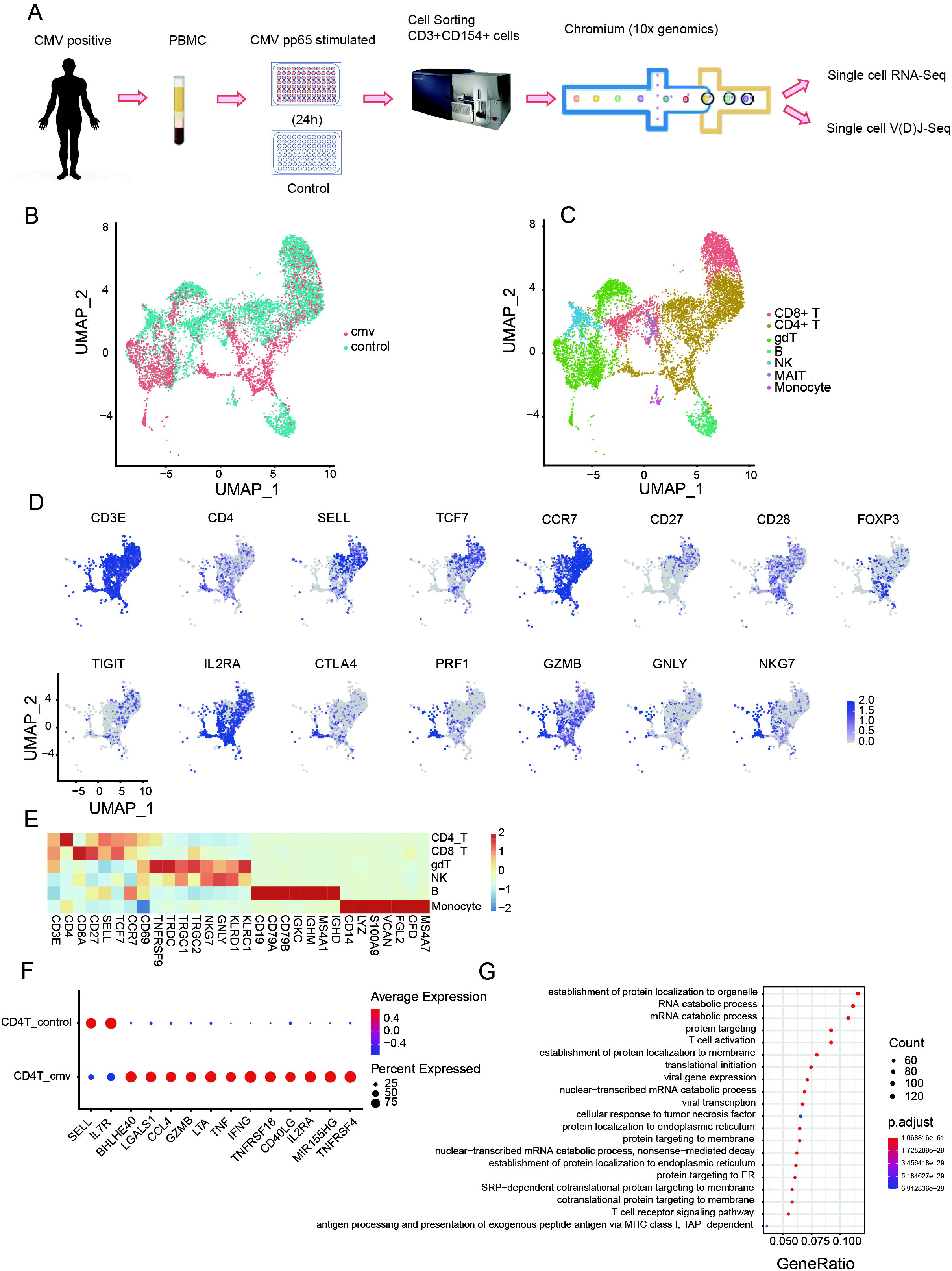
CMV pp65-specific CD4+ T cells are characterized by typical antiviral profiles. (A) Experimental workflow for single-cell analysis of CD4+ T cells from PBMC of three donors includes CMV pp65 in vitro stimulation and culture, CD154+ T cell sorting, and 5′ single-cell RNA and paired T cell receptor sequencing. UMAP embeddings of merged scRNA-seq profiles from control and stimulated (CMV) immune cells were plotted and colored by cell cluster (B) and sample (C) respectively. (D) UMAP projections for the merged CD4+ T cells colored by expression of *CD3E, CD4, SELL* (naive marker), *TCF7, CCR7, CD27, CD28, FOXP3* (Treg marker), *TIGIT, IL2RA, CTLA4, PRF1* (cytotoxic marker), *GZMB, GNLY, NKG7*. Expression values are normalized across CMV and control datasets. (E) Heat map of scaled mean gene expression of the major canonical markers (columns) detected in different cell types in merged cells of CMV and control (rows). (F) Dot plot of differential express genes (DEGs) shows both the expression level and the percentage of CD4+ T cells in CMV and control. (G) Gene ontology (GO) analysis of DEGs between CMV CD4+ T and that of control. The Top 20 enriched GO terms are ordered on the y-axis. X-axis represents the gene percentage in enriched GO terms. Sizes of the dots represent the number of genes included in each GO term. The color gradient of dots represents the adjusted P-values of each enriched GO term.

After stringent quality control and filtering by multiple criteria, RNA-seq data were obtained from 2847 and 6493 single cells from the two libraries (CMV and control), detecting a mean of 3041 and 1947 genes per cell, respectively. Productive VDJ sequences were obtained for 1271 CMV cells and 3557 control cells. The unsupervised clustering of all cells from CMV and control samples suggested that there are multiple subsets of CMV CD4T cells (Figure1. B, C, D). CMV stimulated CD4T cells and control CD4T cells that have both mRNA and VDJ data (CMV: 1200 cells, control: 1911 cells) were used for further analysis.

To reveal the potential function of overall CMV stimulated CD4+ T cells, we analyzed differentially expressed genes between CMV CD4T and control CD4T cells. CMV CD4+ T cells show a typical T cell activation profile including increased expression of *IL2RA, OX40 (TNFRSF4*), *MIR155HG, TNFRSF18, CD40LG* and *LGALS1*, and decreased expression of *IL7R* and *SELL*. These cells also express genes of inflammatory cytokines *IFNG* and *TNF*^29,30^, T-bet independent IFN-γ production inducer *BHLHE40*^31^, pro-inflammatory chemokine *CCL4*, and cytotoxic molecules *LTA* and *GZMB* (Figure 1E). These results suggest that CMV CD4T cells are consist of several groups of activated multiple-cytokine-producing antiviral cells. This result is further confirmed by Gene Ontology (GO) analysis, where DEGs are significantly enriched in pathways such as T cell activation and response to tumor necrosis factors (Figure 1F). Consistent with previous reports using CD154 as a marker for antigen-specific CD4T cells, the cells we obtained here using the same strategy exhibit a typical activated anti-viral response.

### Polyfunctionality profiles of CMV pp65-specific CD4 + T cell subsets

Subpopulations of CD4+ T cell were further identified with canonical markers (Supplementary Table2) and projected onto the UMAP embeddings (Figure 2A,2B). Stimulated CD4+ T cells were clustered into five groups: naïve (17.01% of total CMV CD4+ T cells), memory-like (11.20%), Treg (56.68%), effector memory (Tem, 3.65%), CTL (11.45%). Unstimulated CD4+ T cells were clustered into four subgroups: naïve (55.30%), memory-like (41.98%), Treg (1.78%), and CTL (0.94%) (Figure 2C). The ratio of unstimulated naïve and memory CD4+ T cells is consistent with previous FACS date^32^, implying the classification using the scRNA-seq data in this study has a good consistence with that of FACS. Obviously, Treg, CTL, and Tem are significantly enriched in CMV, and Treg is the largest subset. This distribution is further identified in cells from each donor, although cells from donor 3 were limited (1.58% of total CD4+T cells) (Supplement Figure 1 A, 1B).

**Figure 2:**
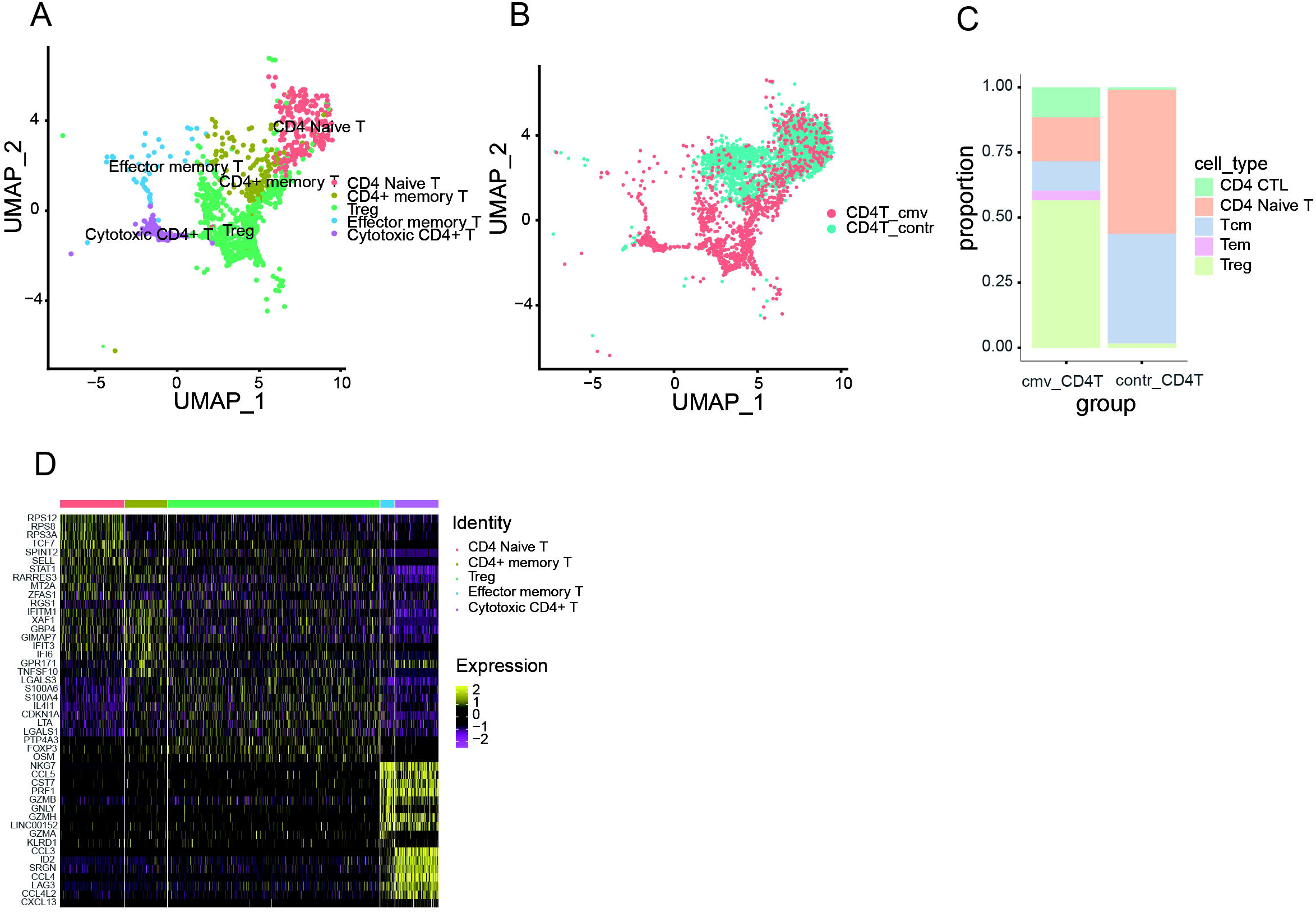
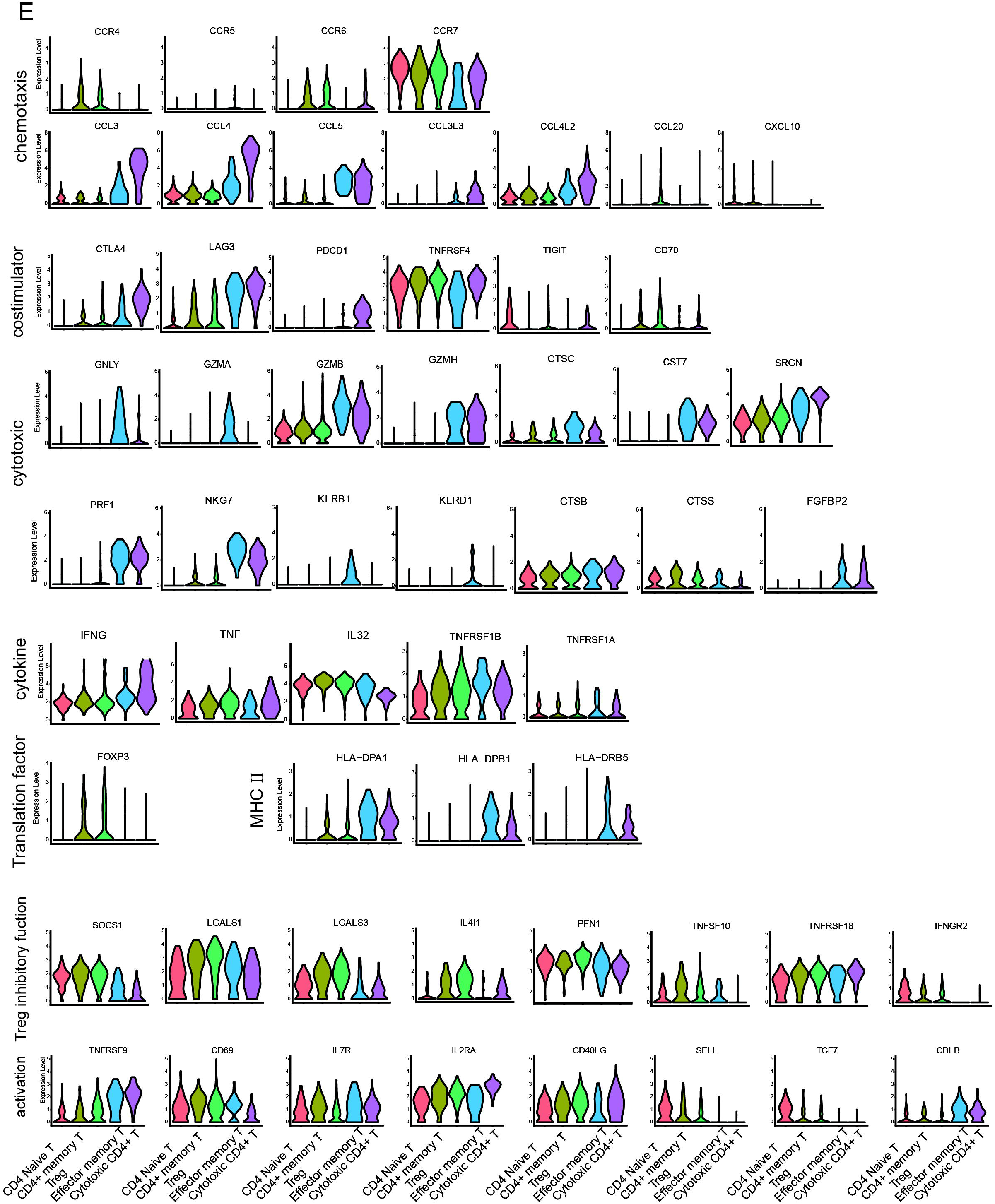
Polyfunctionality profiles of CMV pp65-stimulated CD4 + T cell subsets. UMAP embeddings of merged scRNA-seq profiles from control and stimulated (CMV) CD4+ T cells were plotted and colored by cell cluster (A) and sample (B) respectively. Subpopulations of CD4+ T cell colored in (A) were identified with canonical markers described in Supplementary Table 2. (C) Distribution of the abundance of the five subsets in the CD4+ T cells of CMV and control datasets. (D) Heat map of these five subsets with the Top10 DEGs between each of them. (E) Violin plots of exemplary feature gene expressions of the five subsets. These feature genes were classified and labeled with their group name on the left.

To investigate features of the five CD4+T cell subsets in CMV, we compared them with each other using the *FindAllMarkers* function. The Top10 DGEs were found to be different from each other, indicating that these subsets may have distinct phenotypes (Figure 2D). Therefore, together with the Top 10 DGEs and feature genes from literatures, we analyzed the phenotype of each subset. We firstly analyzed the largest subset, Treg. These cells are *FOXP3*+*IFNG*+*TNF*+, and highly expressed stable marker *SOCS1*, cytotoxic molecules (*LTA, LTB*), and a series of genes relate to inhibition, such as *LGALS1*^33^, *LGALS3*^34^, *IL4I1*^35^ and costimulator *CD70* (Figure 2E). Their stability related cytokine receptors ^36^ (such as *IL2RA, TNFRSF18* [*GITR*], *IFNGR2*), and inhibitory function related costimulatory molecules (such as *LAG3, CTLA4, TNFRSF4* [*OX40*], *TIGIT*), are also abundantly expressed. The expression of chemokine receptor (*CCR4, CCR6, CCR7*) indicate their chemotaxis toward *CCL3, CCL5* (which is highly expressed by CTL and Tem in our data) and homing ability to secondary lymphoid organs of Treg cells, the high expression of *CCR6* and *CCL20* suggests that they can cluster in a self-sustaining positive feedback loop.

Then, we analyzed CD4 Tem and CTL, since they exhibit similar expression profiles (Figure 2E). Tem and CTL both expressed a high level of cytotoxic relative molecules (*GNLY, GZMB, GZMH, CTSC, CST7, PRF1, NKG7, CTSB, FGFBP2*). Those similar expression of cytotoxic markers in CTL and Tem indicate they may employ the same mechanism - the granule exocytosis pathway, to initiate target cell apoptosis. This mechanism involves the regulated release of the contents of cytotoxic granules (such as *PRF1, GZMB, GZMH, GZMA, CTSC, GNLY*), into the immunological synapse formed between the effector and target cell and kill them^37^. Besides, Tem expressed more cytotoxic genes such as *LTB, GZMA, KLRB1*, and *KLRD1* than CTL, indicating the functional spectrum of CD4 Tem is wider than that of CTL. CTL and Tem also abundantly expressed chemokine genes (*CCL3, CCL4, CCL5, CCL3L3, CCL4L2*), MHC □ (*HLA-DPA1, HLA-DPB1, HLA-DRB5*) and co-stimulators (*LAG3, CTLA4, OX40, PDCD1*), indicating they may attract common targets to the inflammatory site and may kill them in another mechanism - MHC class II-dependent fashion^38,39^. Previous studies reported that CTL may stem from Tem^23^. Subsequent TCR repertoire analysis in our data also suggests that CTL may be originated from Tem.

CMV pp65 peptides pool exposed CD4+ naïve T cells in our data show obvious activation characteristics. In total, 981 genes were differentially expressed (adjusted p< 0.05) upon stimulation with the CMV peptides relative to control (Figure 3A; Supplement Table 3), of which 124 and 36 were upregulated and downregulated with a log2-fold change > 1 respectively. These 124 up-regulated genes comprised a group for encoding the cytokines and chemokines (*LTA, MIF, IL32, CXCL10* and *CCL4L2*), a group involving in metabolic [e.g., *GAPDH, PKM, ENO1, TPI1, PGK1*] (Figure 3E) ^40,41^, and a group regulating protein synthesis (e.g., *WARS, SEC61G, EIF5A*). These phenomena support the cell activation^42^. We also found that naïve T cells increased the expression of calcium binding proteins encoded by S100 family genes(e.g., *S100A4, S100A11* and *S100A10*) and cytoskeleton related protein(e.g., *ACTG1, ACTB, TUBB, PFN1, MYO1G*), which were reported in response to TCR engagement by antigen^43,44^. Besides, many regulatory markers (e.g., *GITR* [*TNFRSF18*], *CISH, SOCS1, TIGIT*) and cell apoptosis regulation markers (e.g., *LGALS1, FAM162A, CFLAR, FAS, CDKN1A*) were strongly upregulated to maintain immune balance^45,46^, although their expression level differs in cells at different differentiation period ^47,46,47^. These 36 down-regulated genes including *CD127, CD27* and *CD69L*, and were consistent with previous studies upon T cells activation^48^.

**Figure 3:**
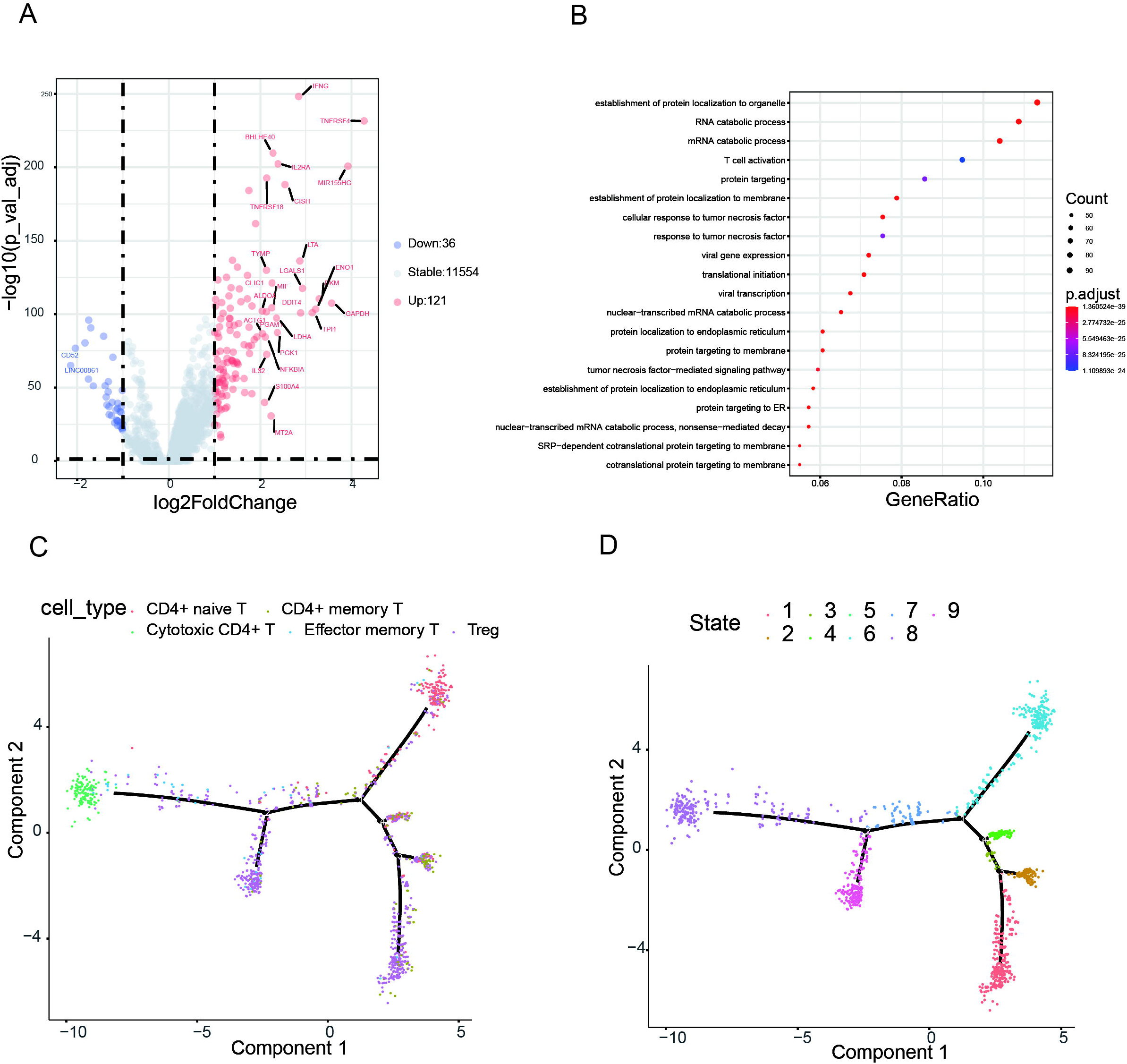
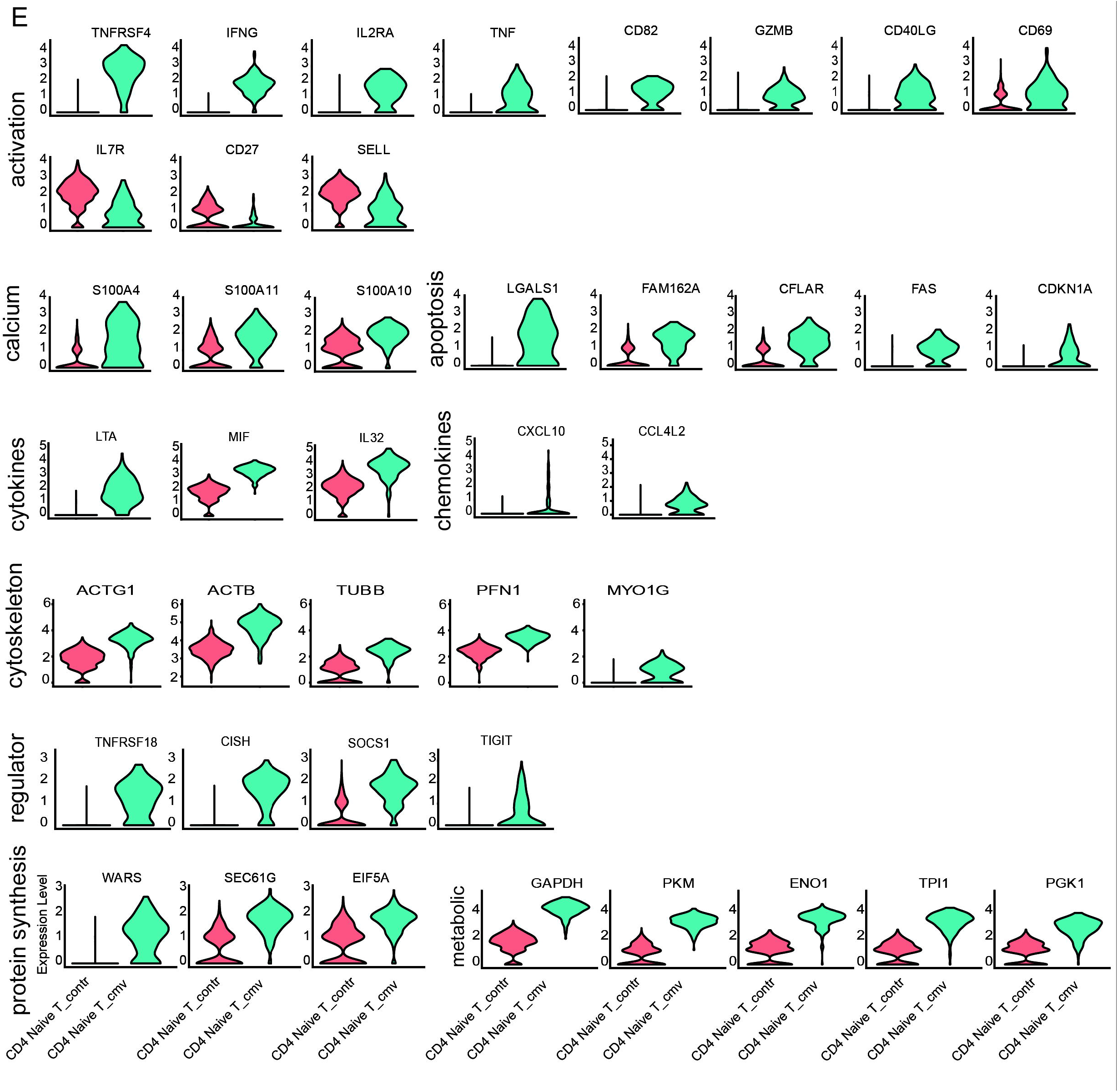
CMV pp65 stimulated CD4+ naïve T cells show obvious activation characteristics. (A) Volcano plot showing the log2FoldChange(x-axis) and -log10(p_val_adj) (y-axis) for differential expression gene between CMV and control naïve CD4+ T cells. Genes with log2FoldChange > 1 and adjusted p value < 0.05 are more highly expressed among CMV and highlighted in red and labeled with their names. Genes with log2FoldChange <-1 and adjusted p value < 0.05 are downregulated in CMV and highlighted in blue and labeled with their names. (B) GO analysis of DEGs between CMV naïve CD4+ T and that of control. The Top 20 enriched GO terms are ordered on the y-axis. X-axis represents the gene percentage in enriched GO terms. Sizes of the dots represent the number of genes included in each GO term. The color gradient of dots represents the adjusted P-values of each enriched GO term. (C) and (D) Pseudotime analysis of the five CD4+ T cell subsets in CMV. These cells were colored by cell cluster (C) and cell state (D) respectively. (E) Violin plots of exemplary feature gene expressions of the naïve CD4+ T cells from CMV and control. These feature genes were classified and labeled with their group name on the left.

GO analysis of the differentially expressed genes between cells from CMV pp56-stimulated and control naïve cells demonstrated the significant enrichment of genes associated with T cell activation, protein processing, viral gene expression, cytokine signaling, and RNA processing ((Figure 3B). Pseudotime Analysis further indicated that CMV CD4+ naïve T cells may differentiate into cytotoxic, regulatory, or other effector helper T cells, which may be a reservoir of CD4+ effector T cells in response to CMV antigen stimulation (Figure 3C, 3D).

### CMV pp65-specific CD4-CTLs shared antigen specificity with Tem

T cell receptor (TCR) repertoire reflects antigen-specificity for cells, and their antigen experience in effector and memory subsets. We therefore analyzed the features of TCR repertoire in each cell cluster from each donor to uncover their potential antigen specificity (donor 1, donor 2). The gene combinations used by each cluster was ranked, showing an obvious clonality of genes combination in CD4-CTL, and a slight genes expansion in Tem. The expanded gene combinations in CD4-CTL and Tem are different between two donors, where Donor1 preferred to use *TRAV5_TRAJ44_TRBV4-3_TRBJ2-1* in CD4-CTL and *TRAV1-2_TRAJ36_TRBV28_TRBJ1-2* in Tem, but Donor2 used *TRAV41_TRAJ49_TRBV6-5_TRBJ2-1* in both CD4-CTL and Tem(Figure 4A). The analyses on CDR3 from TCR alpha chain (TRA) and TRB further verified the clonality in CD4-CTL and Tem. Two expanded clones in CD4-CTL and one in Tem were observed in Donor1, and four in CD4-CTL and one in Tem was observed in Donor2 (Figure 4B). The abnormal features were also exhibited in CDR3 length in naïve, CD4-CTL and Tem, where CDR3 in TRA and TRB from these subsets are shorten in CMV comparing to those in control (Figure 4C). These imply that the repertoire of CD4-CTL and Tem may be largely skewed. To identify clones targeting the same antigens among cell subsets, we used GLIPH2^23^ to cluster clones from CMV and control. We identified convergences between memory and Treg cells, and those between Tem and CTLs in CMV. Interestingly, the convergent TCR clone in CD4-CTL from CMV just composed a minor part of CTL from control in Donor 1, and no convergent clone was found between them in donor 2, suggesting that CMV pp65-reactivated CTLs is rare in total CTLs. Furthermore, the dominated clones in CTLs almost derived from Tem cells in Donor 2, while in donor 1, only a small part of CTLs clones was shared with Tem (Figure 4D). This phenomenon indicates that Tem and CD4-CTL cells shared antigen specificity, and Tem may have a tight relationship with CD4-CTL in differentiation. Meanwhile, we did not find either expanded gene combination nor expanded CDR3 clones in Treg, implying Treg may be unspecific for antigens and activated in TCR-independent manners.

**Figure 4:**
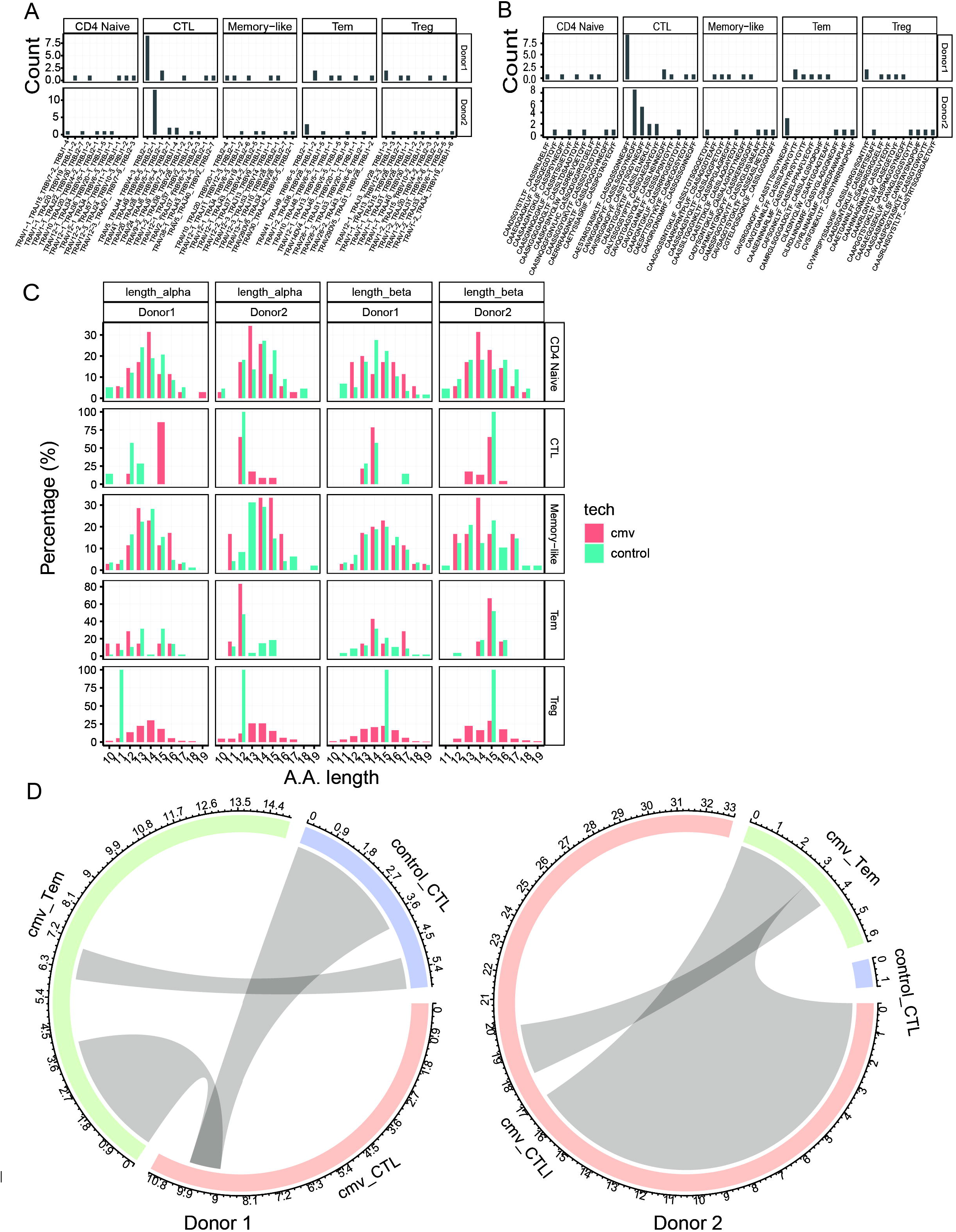
TCR repertoire analysis of the five CD4+ T cell subsets from donor 1 and donor 2. Frequencies of gene combination of TRAV_TRAJ_TRBV_TRDJ (A) and the combination of CDR3a_CDR3b (B) of five stimulated CD4+ T cell subsets from each donor. X-axis is the ranked top 10 gene combination of each cluster. Y-axis is the count of each clonotype. (C) Distribution of amino acid (A.A.) length of TCR alpha and TCR beta of five CD4+ T cell subsets in CMV and control from the two donors. (D) TCR convergence of each cluster from two donors were analyzed with GLIPH2^23^ (left: donor 1, right: donor 2). Clonotypes are sorted by descending order with their sequence counts, and the wide on circus indicate the sequence count per clonotype.

## Discussion

CD154 is an effective marker to combine with single-cell mRNA sequencing and accomplishment high-throughput analysis of virus antigen-specific T cells cytometry^24–27^. Traditional research methods based on secreted cytokines, such as IFNG or TNF, to testing cmv-specific T cells, proven to perform well^49–51^. However, it was of limited application in combination with sc-mRNA sequencing for a reason of cell damage caused by intracellular staining. Another popular method uses Peptide-MHC multimer to isolate antigen-specific T cells according to specific binding of TCR with pMHC allows detailed TCR and phenotypic analysis of cells using single-cell technologies ^52,53,54^. However, the decrease of TCR expression in activated T cells leads to the preference of relative low antigen-specific T cells bound with tetramer, and the selection of multimer-binding CD4+ T cells may bias our understanding of the phenotype of antigen-specific CD4+ T cells^55^. The fact that 83.8% CD4+ T cells in CMV high express *CD154* (*CD40LG*) compared with 17.4% low expressed in control indicate that CD154 is comparable with IFNG and TNF in discriminating antigen-specific CD4+ T cells (Figure 2c). Thus, CD154 is an effective marker to combine with single-cell mRNA sequencing and accomplishment a high-throughput technique platform for research on antigen activated CD4+ T cells.

Bystander activation of CD4+ T cells is less well studied compared with CD8+T cells, but it was demonstrated that unrelated memory CD4 T cells can be activated after a recall tetanus vaccination via bystander activation^56^ and multiple cytokines sharing the common receptor gamma chain can induce CD154/CD40 ligand expression by human CD4+ T lymphocytes via a cyclosporin A-resistant pathway^57^. In our data, CD4+ memory-like T cells are under an environment of IFNG and IL2, and they are prone to be activated by these cytokines. Besides, we find neither clonal expansion nor CDR3 length change in CD4+ memory-like T cell. So, it is difficult to conclude whether these CD4+ memory-like T cells are CMV pp65 antigen-specific.

Our data indicated that the CMV-reactivated Treg had heterogeneously inhibitory functions. *LAG3* and *CTLA4* are classical markers of Treg, by which Treg completely contact MHC-II and CD80/CD86 respectively to repress the activation of conventional T cells. Perforin/granzyme-induced apoptosis is a main pathway used by cytolytic cells to kill target cells^58,59^, and they commonly expressed simultaneously. In our study, Treg highly expressed *GZMB, SRGN* (encoding an element protein for maintain granzyme storage), but their *PRF1* expression is limited. One explanation is that a few perforins is enough to facilitate the entrance of granzyme into target cells. The other is that granzyme B can induce cell death in a perforin-independent manner^60^ where it mediate a cleavage of extracellular matrix to reduce the adhesion of immune cells and result in their death. In addition to these classical inhibitory manners, we observed the expression of *LGALS1* and *LGALS3*, encoding Gal-1 and Gal-3 respectively, which may also participate in Treg immunosuppressive activity^61^. In previous studies, disruption of Gal-1 attenuated the immunoexpressing effect of Treg cells^62^ and Gal-1 from Treg induced the dysfunction of effector T cells and modulated their transient calcium influx^63^. This regulatory mechanism is not limited to Gal-1 but also employed by Gal-3 in Treg^64^.

The presentation of Th1-related cytokines in Treg has been observed in some studies, but whether its appearance implied the stability of Treg inhibitory function is still controversial. In autoimmune diseases, the expression of effector cell cytokines often coupled with decreased inhibitory function in Treg^65,66^; while in other conditions, the expression of T-bet and inflammatory cytokines does not affect the inhibitory function^67,68^. TIGIT is a key regulator to maintain the stability of Treg inhibitory function^69,70^, and is commonly expressed in activated naïve, memory and Treg cells^71^. In this study, Treg expressed TIGIT as similar as activated naïve T cells, implying Treg retained the inhibitory function under CMV infections, and expression of IFN-γ and TNF might enhance their inhibitory function on cells with Th1 phenotypes, including CD4-CTL, Tem and activated naïve T cells. Furthermore, *CCR6, CCR5, CCR7, CCL20, CCL3, CCL4* and *CCL5* were expressed by both Treg^72^ and conventional cells, ensuring co-localization of these cells. Meanwhile, Treg unregulated *IL2RA*, implying an enhanced IL-2 signaling pathway. LTA is the downstream protein of IL-2RA, and its expression condition lymphatic endothelia for enhanced Treg transendothelial migration^73^. Notably, no convergent TCR clone was found between Treg and other CMV-reactivated CD4+ T cells, suggesting Treg may not function to inhibit other CMV-related CD4+ T cells in TCR-dependent manner. In together, Treg cells in CMV infection maintain their inhibitory function, and obtained reinforced inhibitory ability by enhanced migration function.

Another significance of Treg is its large proportion in CD4+ T cells. CD4-CTL in CMV is reported to increase cardiovascular mortality risk of CMV carriers^74^. Chronic infection and repeated activation of CMV may induced a continuous presentation of CD4-CTL and activation of other immune cells, and it is reasonable to induce an accumulation of regulatory cells to avoid their side effects. The large percentage of Treg in CMV-reactivated cells partly supported the hypothesis, but more studies are need to clarify whether the accumulation is necessary to protect individual or contributes to the persistent CMV infection. Additionally, it is novel to find the expression of *CD70* on Treg to our knowledge. *CD70* is commonly expressed on antigen-presenting cells as well as activated T cells to conform CD27–CD70 pathway to provide a costimulatory signal. In T cells, CD70 was showed to induce caspase-dependent apoptosis, and it may also perform a similar function in Treg^75^. However, more investigations should be taken to unveil CD70 function in Treg.

CMV-specific CD4+ Tem displays a distinct cytotoxic function as highly expressed a panel of canonical cytolytic molecules (*GZMB, GZMH, GZMA, PRF1, GNLY, NKG7, IFNG, CTSC, FGFBP2*, and *KLRB1*). GZMA, GZMB, and GZMH belong to granzyme, a subfamily of serine proteases that are function in mediated cell death. This simultaneous high expression of granzyme, PRF1, CTSC, and GNLY, may imply that CMV-specific CD4+ Tem is capable of kill target cells in a granzyme- and perforin-dependent manner. CMV-specific CD4+ Tem cells exert their function in peripheral target organs by the production of antimicrobial lymphokines, Inflammatory chemokine, and cytotoxin, thereby directly contributing to the containment of viral infection. These well prepared cytolytic particles enable CD4+ Tem to quickly start a war against the infection virus.

CD4+ T cells are central organizers in immune responses. CMV-stimulated CD4+ T cell secrets chemokines and effector molecules to recruit and activate or assist other immune cells to orchestral an antiviral response. For example, HCMV-specific CD4+ T cells overall express high levels of chemokines *CCL4* and the CMV-specific naïve CD4+ T cells express high levels of *CXCL10*. CCL4 help recruit immune cells such as macrophages, NK cells, monocytes as well as activated T cells to the site of infection^76–79^. CXCL10 help HCMV-specific CD4+ T to recruit HCMV-specific CD8+ T cells who high expressed CXCR3 (chemokine receptors of CXCL10)^80^ to the Inflammatory site. In addition, IFNG high expressed by CMV-specific CD4+ T could help in direct anti-viral activity and plays an important role in activating immune cells (such as B cells, T cells, NK cells, and macrophage) and augmenting antigen presentation (such as DC, B cells, and macrophages)^81,82^. The high expressed cytotoxic marker (*LTA, GZMB*) also implies possible direct antiviral effects of CD4+ T Cells. Since HCMV has acquired extensive mechanisms of immune evasion, which include the downregulation of MHC class I molecules on infected cells^83,84^. whereas, effector CD4+ T cells require no MHC class I molecules to eliminate target cells, these effector lymphocytes may have evolved to support effector CD8+ T cells in suppression of HCMV infection^1^. Notably, in our data, almost all (99.1%) CMV-specific CD4+ T cell highly expressed IFNG compare with few (1.6%) expressed in control CD4+ T with logFC = 4.576. These results go nicely in line with the fact that CD4 T cells are important in maintain CD8 T-cell activity during prolonged infection, and their role in helping the antiviral antibody response may also be essential^82^.

Unlike other latent-virus specific T cells (such as EBV, HSV), CMV CD4+ T cell does not show an exhaustion phenotype. Both EBV-specific CD8+ and CD4+ T cells are highly susceptible to apoptosis, while CMV CD4+ T cells are not. Akin to the cmv CD4+ T cell response, the majority of activated EBV-specific CD4+ T cells exhibit Th1-type response and express both perforin, granzyme B, CD107a^85,86^ and lack the lymphoid homing markers CCR7 and CD62L^87^. It was speculated that EBV-specific CD4+ T cells might highly effective against MHC-II positive EBV infected B cells. CD4+CD25+ Treg cells are also found in HSV-1 specific CD4+ T cells^88^. HSV^89^: HSV-1 can establish a latent infection in TG of host, while CD4+ and CD8+ T cells can control the reactivation of the HSV-1 by surrounding latently infected neurons. CMV CD4+ T cells exhibit downregulation expression of IL7R, which is also observed in other chronic infections such as LCMV clone 13 and HIV infection^90–93^. This phenomenon is consistent with the previous reports that T cells fail to re-express the IL7R^94^, when they are continuously stimulated.

## Methods and Materials

### PBMC preparation

We obtained peripheral blood from three CMV IgG-positive, heathy donors though a research protocol proved by the Beijing genomics institution-Shenzhen (BGI-Shenzhen) Institutional Review Board (IRB). Peripheral blood mononuclear cell (PBMCs) were immediately isolated from blood collected with EDTA blood collection tube by density centrifuge method with Histopaque-1077 (Sigma, Cat. 10771) within two hours^95^, resuspended in 4°C cryopreservation medium consisting 90% fetal bovine serum (FBS, HYCLONE, Cat. sh30084.03) and 10% Dimethyl sulfoxide (DMSO, Sigma, Cat. D4540) and then placed in Mr. Frosty (Thermo Scientific) in −80°C container. Samples were then moved to liquid nitrogen for a long-time storage.

Additionally, 2 ml peripheral blood from each donor was collected by blood collection tube without any additive, placed at room temperature for 30 minutes(min) and centrifuged for 10min at 2000g. Then plasma was collected and heat shocked for 30min at 55°C.

### PBMC stimulation

Frozen PBMC from liquid nitrogen were immediately thawed in 37°C water and resuspended in complete medium (RPMI 1640 medium, 10% NEAA and 2% autologous plasma; RPMI 1640 and NEAA were purchased from ThermoFisher with Cat. 72400120 and Cat. 11140050) to a final density of 1*10^7 per milliliter (ml). We moved 150 microliter (ul) of cell suspension with three repetitions to each well in the 96-well U-plate (Falcon) and incubated them at 37°C for two hours. Then 75ul culture supernatant in each well was replaced by 75ul stimulation medium, and gently mixed. Cells were cultured in an incubator with 5% CO_2_ at 37°C for 24 hours.

The stimulation medium included RPMI 1640 medium (without serum), anti-CD28 (2ug/ml, Clone G28.5, Genetex, Cat. GTX14148), anti-CD40 (2ug/ml, Clone HB14, Miltenyi, Cat. 130-094-133) with/without CMV peptide (1.2 nmol/ml per peptide). To preserve the surface expression of CD154 on activated T cells, we used anti-CD40 to inhibit the interaction of surface CD154 with its counterpart CD40 as described in the previous study^22^. CMV pp65 peptide was purchased from Miltenyi (Cat. 130-093-438) and diluted in sterile water.

### Enrichment of CMV pp65-specific T cells

Cells were collected and washed with FACS washing buffer (DPBS, 2% FBS and 1mM EDTA) for once and resuspended in staining buffer (FACS washing buffer with 10% human plasma and 1% BSA) containing antibodies against CD3, CD4, CD154 and CD69 (Supplementary Table 1). After been incubated on ice for 40 minutes, cells were washed with FACS washing buffer for twice, and resuspended in 100ul washing buffer. The stained cells were analyzed and sorted by a BD FACS Aria II cell sorter (BD Biosciences). For cells stimulated with CMV peptide, CD3+CD154+ cells were sorted as CMV-specific T cells; For unstimulating cells, monocytes and lymphocytes were sorted respectively and re-mixed as a control. The gating schedule for cells sorting was recorded by BD Aria II (Supplementary Fig.1), and FACS data was analyzed with Flowjo v10.0.7

### Droplet generation, 10X RNA-seq and TCR-seq library preparation and sequencing

After been counted with C-Chip (inCYTO), CMV-specific cells and control cells from all three individuals were mixed separately, diluted with PBS to a final concertation ∼800/ul, and about 20,000 cells per reaction were loaded onto a Chromium Single Cell Chip (10x Genomics). The libraries for RNA-seq and TCR-seq were prepared using the Chromium Single Cell 5’ Library & Gel Bead Kit v2, Chromium Single Cell V(D)J Human T Cell Enrichment Kit (10X Genomics) following the manufactory’s protocol. Sequences within these libraries were ligated with BGIseq adapters ^70^(doi:10.1016/j.scib.2020.01.002), and then CMV and control libraries were loaded onto sequencing chip. The RNA-seq libraries were sequenced with an 8-base index read, a 26-base read 1 containing cell-identifying barcodes and unique molecular identifiers (UMIs), and a 100-base read 2 containing transcript sequences on BGIseq500; TCR-seq were sequenced with an 8-base index read, a 150-base read 1 containing cell-identifying barcodes, UMIs and insert started from V-gene region, and a 150-base read 2 containing insert from C-gene region. The raw data after sequencing was about 10+35 Gb per library for RNA-seq and 35+35 Gb for TCR-seq. The data that support the findings of this study have been deposited into CNGB Sequence Archive^96^ of CNGBdb^97^ with accession number CNP0001262.

### Preprocessing single cell RNA-seq data

Raw data were split according to sample barcodes into CMV-stimulated (ST) and unstimulated library (CON), and then were filtered, blasted, aligned and qualified by Cellranger v2.2.0 with reference of refdata-cellranger-GRCh38-1.2.0 for RNA-seq data and Cellranger v3.0.0 with refdata-cellranger-vdj-GRCh38-alts-ensembl-2.0.0 for TCR-seq data. Other parameters were set as default in the software.

**Supplementary Table 1.**
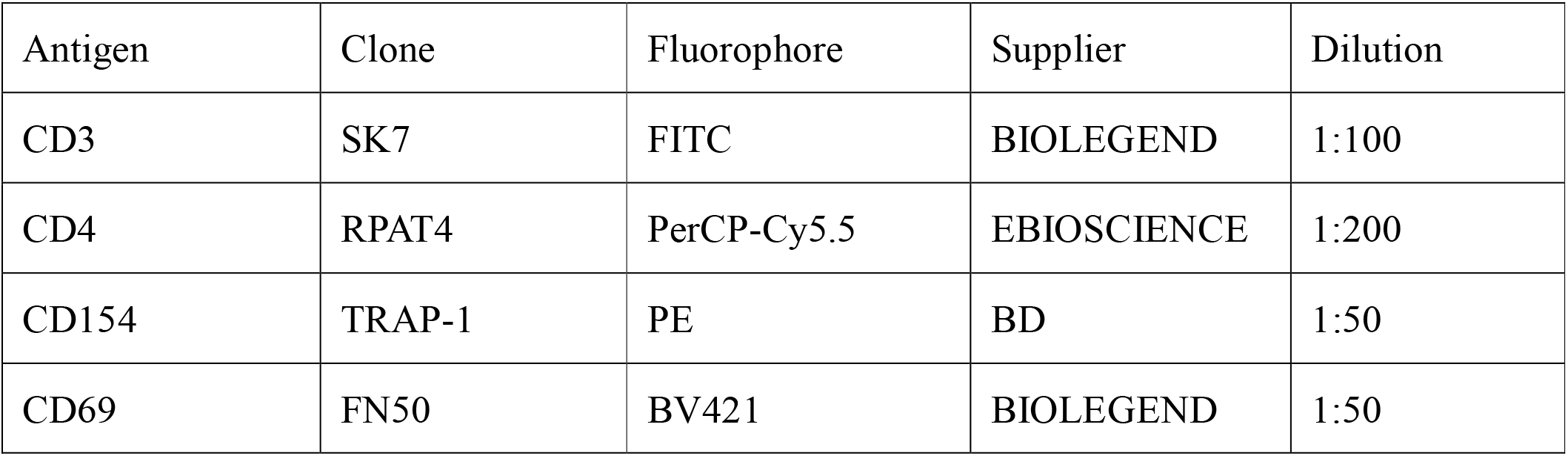
FACS antibodies.

### Data Integrating and cell clustering

The R package Seurat^98^ 3.1.5 was used to integrate and analyze datasets from CMV and control. The merged expression matrix was firstly filtered following the Seurat recommendation^99,100^ and a total of 8671 cells with unique UMI was obtained. Unsupervised clustering was conducted with Seurat with the parameter res = 0.5, it revealed a total of 16 clusters. We used mRNA biomarkers (Supplemental Table 2) obtained from recently published articles^101–103^ to identify these clusters, and GSEA analysis was conducted to identify cluster3(see method Gene set enrichment analysis). TCR repertoire datasets were also combined to identify putative mucosal associated invariant T (MAIT) cells and γδT. Based on the expression of a typical TCR ^104,105^(Supplemental Table 2), a 17th cluster (putative mucosal associated invariant T, MAIT) was emerged from cluster12. Based on whether they express a TCR a chain, and meanwhile express γδ biomarkers (CD3D, CD3E, TRDC, TRGC1, TRGC2), 2111 γδT cells were identified and excluded from further analysis. The above classification was consistent with hematopoietic differentiation and the previously published t-SNE plots of PBMC scRNA-seq^106–108^.

**Supplementary Table 2.** cell type markers.

| Cell Type | Markers |
| --- | --- |
| naïve CD4 T | CD3E, CD4, SELL, CD27, TCF7, CCR7<sup>102</sup> |
| naïve CD8 T | CD3, CD8A, SELL, CD27<sup>102</sup> |
| $\gamma\delta$T | CD3+CD4-CD8- \| CD3+CD4-CD8aa+, CD3D, CD3E, TRDC, TRGC1, TRGC2<sup>101</sup> |
| Memory Like T | CD3E, CD4, SELL, CD27 \| CCR7, SELL, TCF7 |
| Treg | CD3E+, TIGIT, CTLA4<sup>108</sup> \| FOXP3, IL2RA, CTLA4<sup>109</sup> |
| Tem | CD3E, CD4/CD8A, CD27, PRF1, GNLY |
| B | CD79A, CD19, CD79B, IGKC, IGHM, MS4A1, IGHD |
| NK | CD3E-, CD4+, FOXP3+, TNFRSF9+, NKG7, GNLY, NKG7, KLRD1, KLRC1<sup>103</sup> |
| active CD8 T | CD3E, CD8A, CD69, TNFRSF9(CD137) |
| CTL | GZMA, GZMB, PRF1, NKG7,CCR7, CD27, CD28, IL7R<sup>110</sup> |
| Monocyte | LYZ, S100A9, CD14,FGL2,MS4A7<sup>103</sup> |
| MAIT | TRAV1-2 / TRAJ33 , TRAV1-2 / TRAJ20 \| TRAV1-2 / TRAJ12<sup>105</sup> |

### Differential express gene (DEG) analysis

DEG analysis was conducted by the function *FindMarkers* provided by *Seurat*. To character the features of CMV-specific CD4+ T cell response, we used a stricter standard to filter out DEGs between CMV and control CD4+ T cells according to the following standard: for upregulation genes in CMV, adjusted P-value < 0.05, log fold change >1, percentage of cells expressing the gene in CMV sample (pct.1) >0.8, percentage of cells expressing the gene in control (pct.2) < 0.2; for downregulation genes in CMV, adjusted P-value < 0.05, logFC >1, pct.1 <0.2, pct.2 >0.8.

### Quality control metrics and filtering

We used the CellRanger v2.2.0 software with the default settings to process the raw FASTQ files, align the sequencing reads to the GRCh38 transcriptome, and generate a filtered UMI expression profile for each droplet.

### Identifying the sample identity of each droplet

Firstly, transcriptome of each donors’ peripheral blood mononuclear cell was sequenced on the BGI-SEQ500 platform with the sequencing type SE200. Raw data with 10G per sample was obtained. Then, we followed the best practices workflows recommend by GATK(https://gatk.broadinstitute.org/hc/en-us/articles/360035531192-RNAseq-short-variant-discovery-SNPs-Indels-) to call SNP (Single-nucleotide polymorphism) and create VCF files containing the genotype (GT) to assign each barcode to a specific sample. The VCF file and BAM files produced by cellranger2 was passed to the demuxlet software to deconvolute sample identity^28^. Finally, we assign the best guess of the samples’ identity to their corresponding donor and each “possible” or “ambiguous” droplets as unclear.

### Gene ontology analysis

To annotate the potential functions of the DEGs of each CD4+ T cell cluster, GO enrichment analysis was conducted using the clusterProfiler R package^111^(version 3.14.3) with the differential expressed feature genes identified by Seurat. Top 20 Enriched pathways, ranked by normalized enrichment score, with the FDR q-val ≤0. 05 were chosen and visualized.

### Gene set enrichment analysis

Gene set enrichment analysis (GSEA, http://www.broad.mit.edu/gsea) was conducted with default sets to deduce the cell type of cluster 3. The gene set collection used for GSEA was c7.all.v7.1.symbols.gmt (ftp.broadinstitute.org://pub/gsea/gene_sets/c7.all.v7.1.symbols.gmt).

### Pseudotime Analysis

A total of 1200 cells from cmv (include five clusters: naïve CD4 T, central memory T, effector memory T, Treg, CTL) were used for pseudotime analysis. The analysis was performed with Monocle2 (version 2.14.0) with UMI count expression data ^112^. All the parameters were set as default. Differential expression gene between the five clusters was identified with differentialGeneTest function implemented in Monocle 2. The resulting genes with qval < 1e-5 was selected for ordering cells in pseudotime using reduceDimension with the DDRTree method and orderCells functions.

**Donor 1.**
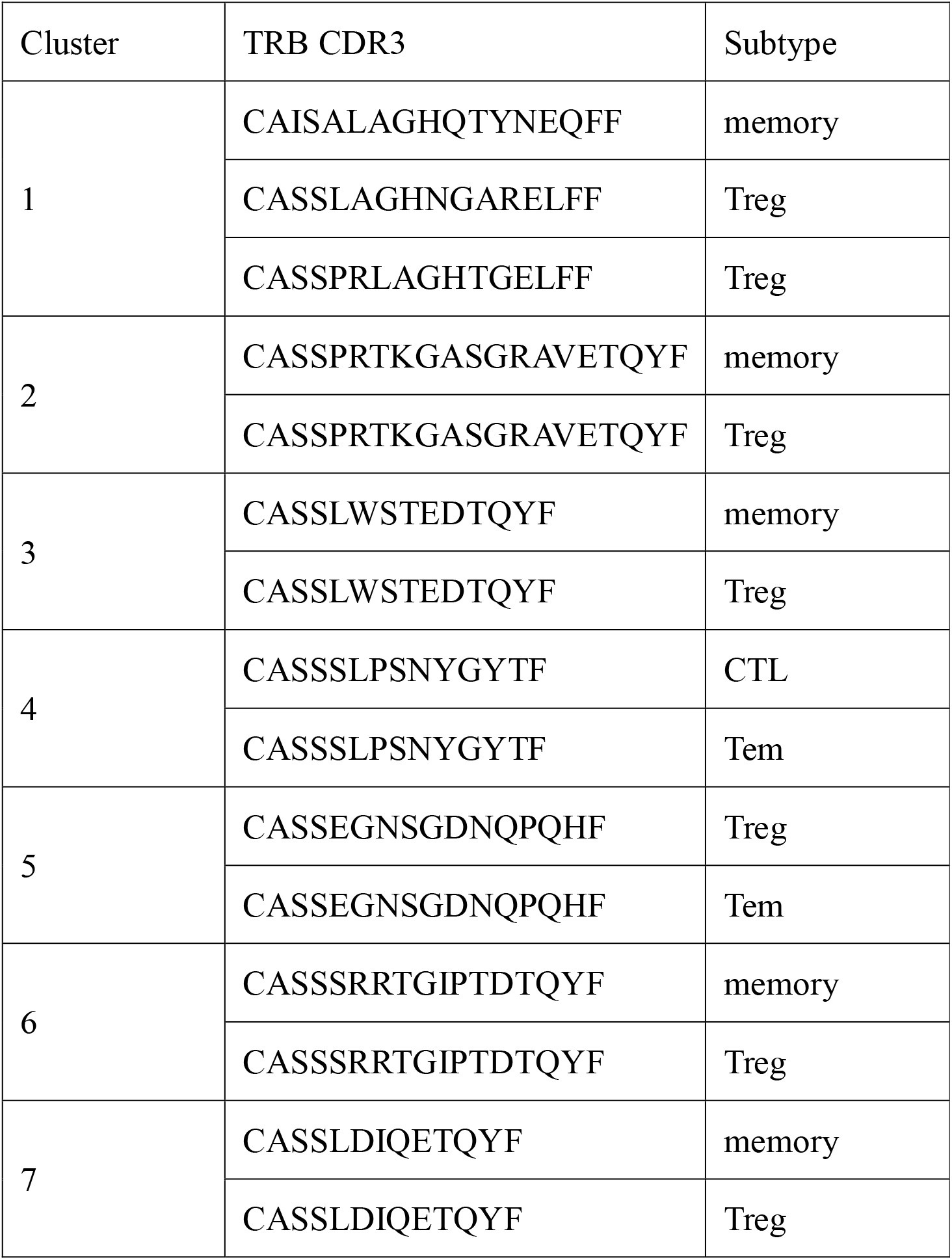

| Cluster | TRB CDR3 | Subtype |
| --- | --- | --- |
| 1 | CAISALAGHQTYNEQFF | memory |
|  | CASSLAGHNGARELFF | Treg |
|  | CASSPRLAGHTGELFF | Treg |
| 2 | CASSPRTKGASGRAVETQYF | memory |
|  | CASSPRTKGASGRAVETQYF | Treg |
| 3 | CASSLWSTEDTQYF | memory |
|  | CASSLWSTEDTQYF | Treg |
| 4 | CASSSLPSNYGYTF | CTL |
|  | CASSSLPSNYGYTF | Tem |
| 5 | CASSEGNSGDNQPQHF | Treg |
|  | CASSEGNSGDNQPQHF | Tem |
| 6 | CASSSRRTGIPTDTQYF | memory |
|  | CASSSRRTGIPTDTQYF | Treg |
| 7 | CASSLDIQETQYF | memory |
|  | CASSLDIQETQYF | Treg |

**Donor 2.**

| Cluster | TRB CDR3 | Subtype |
| --- | --- | --- |
| 1 | CASSQVGAELYGYTF | Treg |
|  | CASSQVGTATEAFF | Treg |
|  | CASSSQVGYGYTF | memory |
| 2 | CASSTTSGGYNEQFF | CTL |
|  | CASSTTSGGYNEQFF | Tem |
| 3 | CSAREDRAWAPLHF | CTL |
|  | CSAREDRAWAPLHF | Tem |

---

Supplement Figure 1: Distribution of stimulated CD4+T cells from each donor (n = 3). (A) UMAP embeddings of CMV CD4+ T cells from each donor. *demuxlet*^28^ was used to assign these cells to each donor, the ambiguous droplet was assigned as “unclear”. Proportions of cells from each donor was showed on the left. The UMAP embeddings were colored (A) or split by donors(B) respectively.

