## Supplementary figures and images for "Dissecting the landscape of activated CMV-stimulated CD4+ T cells in human by linking single-cell RNA-seq with T-cell receptor sequencing"

### Supplemental Figure1

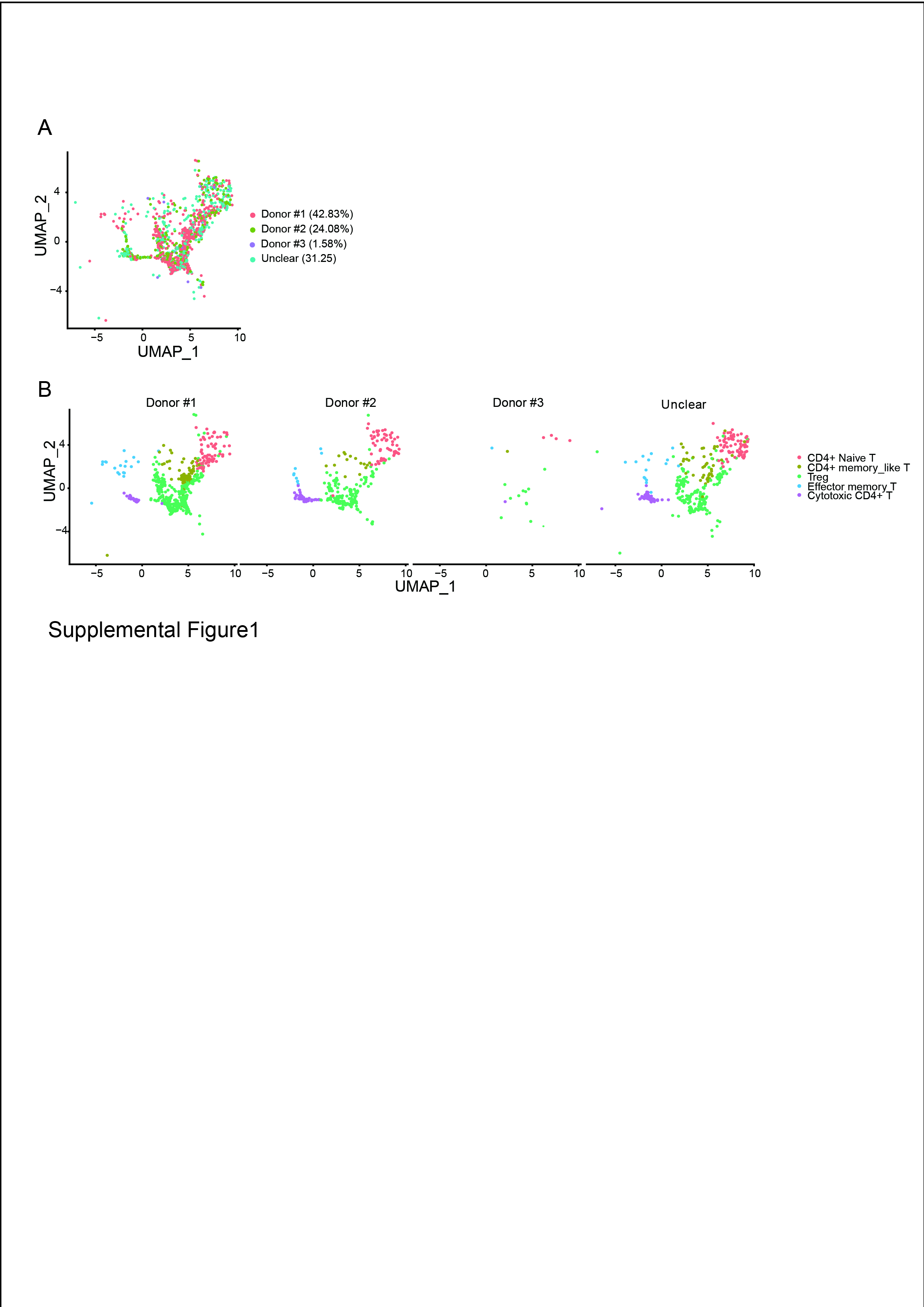
